# A Recessive *oca2* Mutation Underlies Albinism in *Xiphophorus* fish

**DOI:** 10.1101/2025.01.20.633999

**Authors:** Yanting Xing, William Boswell, Jessica Parker, Kang Du, Manfred Schartl, Yuan Lu

## Abstract

Oculocutaneous albinism (OCA) is a group of genetic disorders characterized by impaired melanin production, leading to reduced pigmentation in the skin, hair, and eyes. *Xiphophorus*, a genus of small freshwater fish, has been a pivotal model organism in pigmentation disorder research, providing key findings in the genetic pathways governing physiological and pathological pigment cell biology. Leveraging the well-established research framework provided by *Xiphophorus*, we have identified a spontaneously occurring albinism phenotype in swordtail fish *Xiphophorus hellerii*. Genetic mapping of albino fish showed that albinism is associated with a recessive mutation in the *oca2* gene. This discovery provides a novel opportunity to explore functions of *oca2* gene in pigment cell differentiation, pigment synthesis, melanosome assembly and transportation function and amelanotic melanoma development.

## Introduction

Albinism encompasses a spectrum of inherited disorders characterized by defects in melanin production, resulting in reduced or absent pigmentation in the hair, skin, and eyes^1^. There are 7 types of oculocutaneous albinism (OCA)^2^. They are characterized by distinct phenotypes, ranging from complete lack of pigmentation in OCA1A, caused by mutations in the TYR gene, to milder hypopigmentation and variable vision issues in subtypes such as OCA2 and OCA4, which result from mutations in the OCA2 and SLC45A2 genes, respectively. OCA3, caused by mutations in the TYRP1gene, is often associated with reddish-brown skin and hair, while the rarer subtypes OCA5, OCA6, and OCA7 are associated with intermediate pigmentation and visual impairments. OCA6 is linked to mutations in the SLC24A5 gene, OCA7 is associated with mutations in C10ORF11, and the genetic cause of OCA5, mapped to chromosome 4q24, has yet to be identified. Due to their infrequency, these rarer subtypes exhibit less well-characterized phenotypic variability.^2^ OCA2 is the most prevalent albinism worldwide, with an especially higher incidence for the African Black OCA patients^3,4^. Besides human, OCA is also observed in other vertebrates, including fish^5–7^, pig^8^, rodents^9,10^, water buffalo^11^, and rhesus macaque monkeys^12^, in which the same phenotypes, light body coloration along with pink or red eyes, are shared with mammals.

Beyond its characteristic effects on pigmentation, albinism significantly influences both vision and sleep, as illustrated by studies in *Astyanax mexicanus* cavefish^13^ and rhesus macaques^12^. O’Gorman et al.^13^ demonstrate that OCA2 mutations in cavefish not only lead to albinism but also contribute to reduced sleep duration, a behavioral trait that likely supports increased foraging activity and survival in nutrient-scarce cave environments. In rhesus macaques with albinism^12^, research reveals that mutations in the tyrosinase (TYR) and OCA2 genes result in visual impairments, including foveal hypoplasia and retinal hypopigmentation, providing valuable insights into the genetic and clinical parallels of oculocutaneous albinism in humans. These findings highlight how OCA2 is associated with both visual and behavioral traits, contributing to adaptations necessary for survival in challenging environments.

Additionally, different mutations in the *OCA2* gene among humans are associated with an increased risk of melanoma^14–19^. Utilizing case-control studies, single nucleotide polymorphism (SNP) genotyping and sequencing, hypomorphic allele p.V443I^18,19^, along with p. R305W^14^ and p.R419Q^15,17^, have been significantly associated with increased melanoma risk in specific populations. These OCA2 variants influence melanin production and pigmentary pathways, contributing not only to the pigmentation characteristics of melanomas but also to their risk and pathological diversity^14^. Evidence has been shown melanin-photosensitized radical product is the major causative step of melanoma^20^. This mechanism indicates that variations in melanin types and concentrations, influenced by OCA2 gene variants, enhance radical formation upon UV exposure. Such an increase in radicals can lead to more DNA damage, thereby raising the risk of melanoma development.

*Xiphophorus* is a genus of 26 fresh water small fish species. They are used to study a wide range of diseases etiology or resilience^21^. The *Xiphophorus* species exhibit a wide array of body coloration patterns and are therefore implemented for investigation of genetics underpinning these traits. The best example refers to genetics underlying melanomagenesis^21–24^. However, the understanding melanophore biology, including genetics pathways affecting melanophore differentiation, proliferation, migration, melanosome assembly and intracellular trafficking, are understudied. Therefore, mutants exhibiting pigmentation pattern abnormality will offer unique opportunities to investigate the melanophore biology.

In this study, we identified a spontaneously occurred albinism in swordtail fish *Xiphophorus hellerii* (*X. hellerii*). The albino individuals are characterized by absence of cutaneous and ocular pigmentation. We conducted genetics mapping and identified variant associated to the albino phenotype is a recessive mutation of *oca2* gene.

## Material and Methods

### Research animals

Fish were raised at the fish facilities of the Biocenter of the University of Würzburg and at the *Xiphophorus* Genetics Stock Center at Texas State University. All fish were kept and sample taken in accordance with approved experimental protocols through an authorization (568/300-1870/13) of the Veterinary Office of the District Government of Lower Franconia, Germany, and approved protocol by Texas State University IACUC (IACUC9048). The following laboratory lines were used: *X. hellerii* (strain hIII WLC 1336; origin in Rio Lancetilla, wild type pigmentation pattern). Albino swordtail strain (WLC 2378): the albino mutation from an ornamental swordtail strain was introduced into the genetic background of *X. hellerii* (hIII, WLC 1336) by repeated backcrossing. For mapping the albino locus, WLC 2378 males were crossed to females of strain WLC 534 which is heterozygous for the albino locus and the Spot Dorsal (Sd)-allele of *xmrk* in the genetic background of *X. hellerii* (hIII WLC 1336). Offspring were typed for albino (N=32) or wild type (N=59) pigmentation and absence or presence of *xmrk*.

### DNA and RNA isolation

Fin clips from wild typeand albino *X. hellerii* males were collected and digested with Proteinase K at room temperature for 1 hour. The lysates were transferred to 2.0 mL collection tubes, and DNA was isolated using the QIAcube HT automated bio-sample isolation system (Qiagen, Valencia, CA, USA) with reagents from the QIAamp 96 DNA QIAcube HT Kit. The system features a robotic arm equipped with eight pipettes capable of picking and ejecting pipette tips, self-cleaning, and transferring liquids between wells, columns, or reservoirs in standard 96-well plate formats. Each sample was handled independently throughout the isolation process. DNA concentrations were measured using a Qubit 2.0 fluorometer (Life Technologies, Grand Island, NY, USA) and adjusted for sequencing library preparation.

Skin samples were homogenized in TRI-reagent (Sigma Inc., St. Louis, MO, USA), followed by the addition of 200 μL/mL chloroform, vigorous shaking, and centrifugation at 12,000 g for 5 minutes at 4°C. Total RNA was purified using the RNeasy Mini RNA Isolation Kit (Qiagen, Valencia, CA, USA), with on-column DNase digestion performed at 25°C for 15 minutes to eliminate residual DNA. RNA concentrations were determined using a Qubit 2.0 fluorometer (Life Technologies, Grand Island, NY, USA), and RNA quality was assessed using an Agilent 2100 Bioanalyzer (Agilent Technologies, Santa Clara, CA, USA). Only samples with RIN scores above 7.0 were used for subsequent gene expression profiling.

### Genome wide association study

DNA sequencing reads were mapped to *X. hellerii* genome (GenBank assembly accession: GCA_003331165.2) using Bowtie2^25^. Mpileup files were made using samtools and genotyping was processed using Bcftools^26,27^ Genotype in this study refers to inheritance of wild type or mutant alleles, with heterozygous meaning that a locus exhibited both allele calls, and homozygous means that a locus exhibited only mutant allele calls. Genotypes calls with high statistical confidence were kept for following analyses (i.e., Bcftools: MAPQ□ ≥ □30, Phred score of genotype call□=□0, with alternative genotype call Phred score□ ≥ □20). Per polymorphic locus, a Chisq test was used to test if genotype distribution is biased from an expected 50%-50% distribution of heterozygotes and homozygotes in both wild type and albino population. The Chisq test p-values were adjusted using Bonferroni method to for multiple test correction and were converted to −10*Log10p-value for Manhattan plot.

### Sequence analyses and polymerase chain reaction

RNA sequencing reads were mapped to *X. hellerii* genome (GenBank assembly accession: GCA_003331165.2) using Tophat2.^28^ The short read mapping results in BAM format were indexed and visualized using table genome browser IGV that is configured for *X. hellerii* genome.^26,29^ Each homozygous variant in albino samples were manually analyzed for codon-change mutation. Sashimi plot was made using IGV to show sequencing reads density at exon-exon junctions.

Total RNA was isolated from skin of both wild type and albino fish. Reverse transcription was conducted using total RNA with random primers. To amplify exon 16, primers were designed on *oca2* (RefSeq: NC_045680.1) exon 16. The forward primer sequence is: 5’-AACTGAAGCATGAGATCCTG-3’; The reverse primer sequence is: 5’-CTCTGGAAGGTTTTCATCAT-3’. Electrophoresis was performed to test the PCR amplicon size, followed by amplicons recovered and Sanger Sequencing performed (Azenta Life Sciences, Burlington, MA, USA).

### Protein structure prediction

The amino acid sequence of the *Xiphophorus* wild type *oca2* was retrieved from NCBI (XP_014326265.1). The mutant allele protein sequence was translated from coding sequencing that is constructed by adding sanger sequencing recovered insertion into wild type *oca2* protein sequence. Both wild type and mutant OCA2 protein structures were modeled using AlphaFold3^30^. The amino acid sequences were formatted in FASTA and put into Alphafold3 to generated structural prediction for each isoform. The models with the highest Predicted Template Modeling (pTM) scores were selected for further analysis. Structural alignments and visualizations were conducted using PyMol to assess the impact of the alterations on the protein structure.

## Results

### *X. hellerii* albinism is a monogenic recessive coloration disorder

The *X. hellerii* exhibits three main types of chromatophores: melanophores, which synthesize black or dark-brown eumelanin; xanthophores, which contain carotenoid-based yellow pigments; and erythrophores, which contain pteridine-based red pigments^31^. Additionally, iridophores produce iridescent blue, green, or silver colors through the reflection of light by guanine crystals. The melanophores usually locate between fin rays and at the edge of scales, erythrophores locates around the lateral line, and xanthophores are in the dorsal fin and also caudal fin. The albino *X. hellerii* exhibit no melanophore pigmentation pattern but intact erythrophores and xanthophore pigmentation patterns. In addition, there is no ocular pigmentation and the fish have red eyes (Fig. 1B).

**Figure 1.**
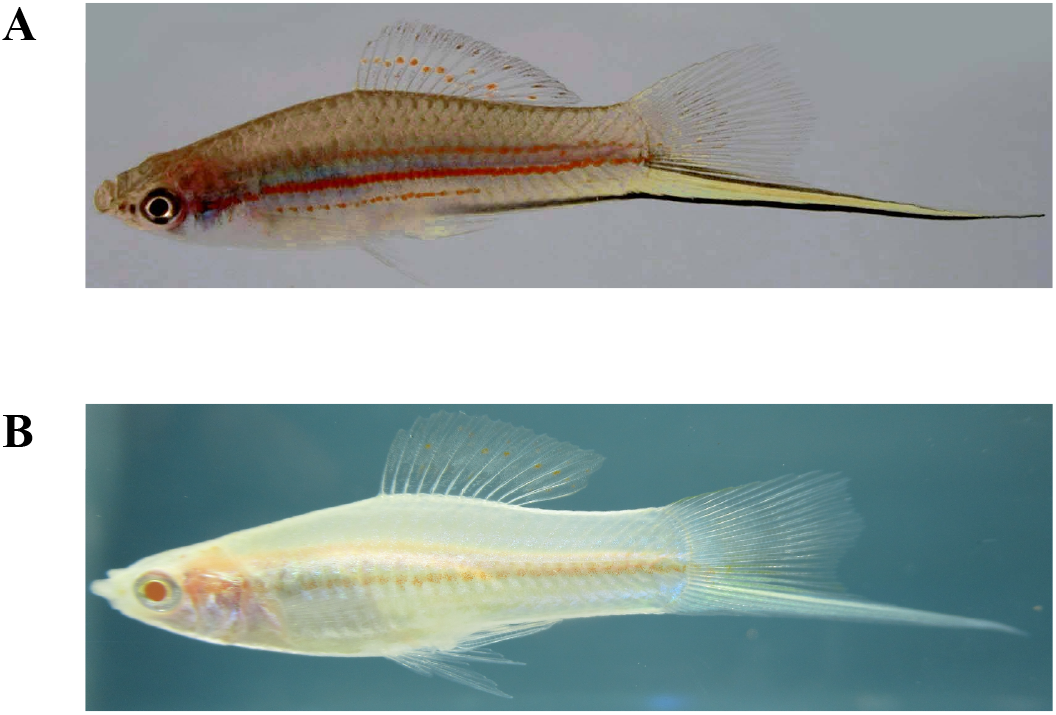
Wild type and albino *X. hellerii*. (A) Wild type *X. hellerii* exhibit macromelanophores (black) in the eyes, at the edge of scale, and between fin rays. Erythrophores (red) appear along the body side lateral line and between fin rays. (B) Albino *X. hellerii* exhibit no macromelanophore pigmentation patterns in the eye, skin and fins.

**Figure 2.**
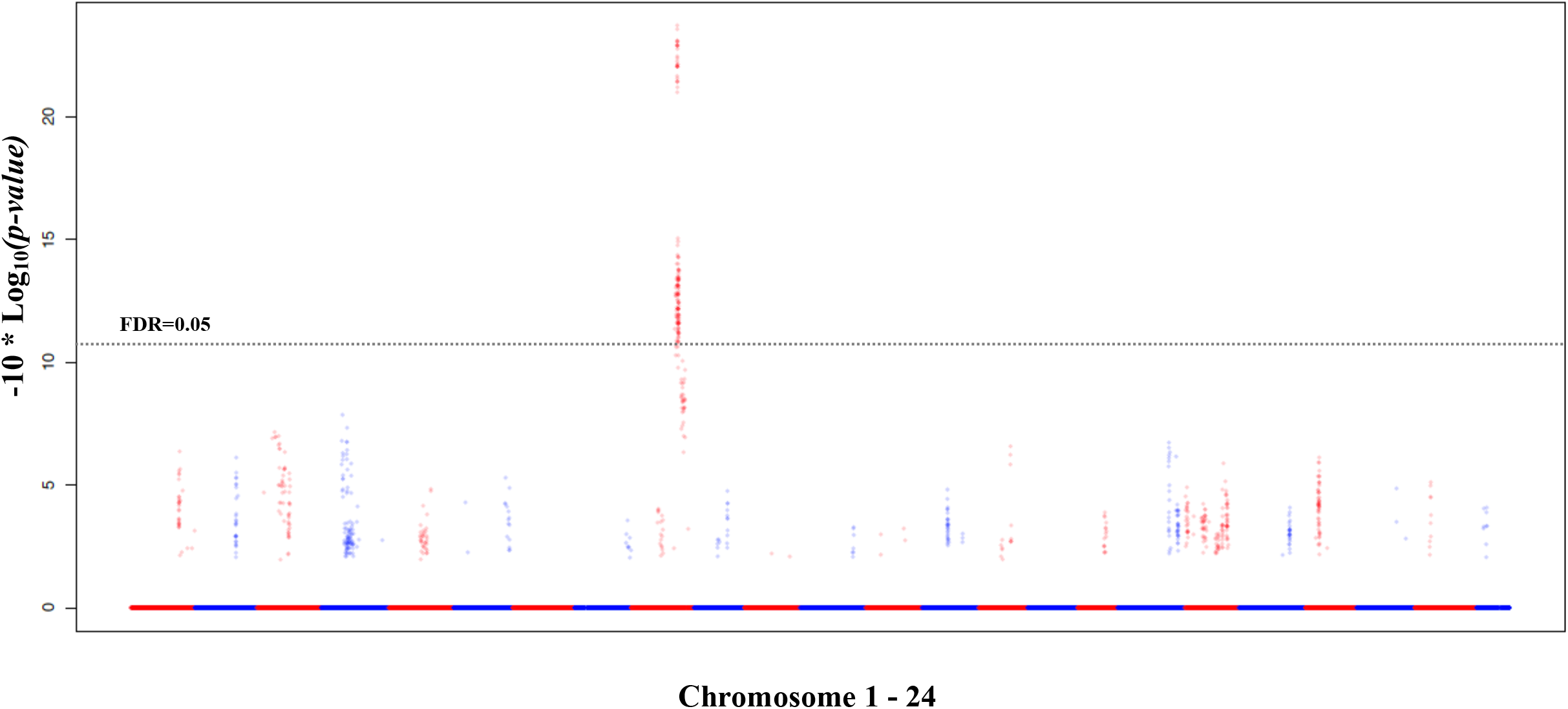
Genetic mapping of genetic locus underlying albinism. Hybrids were produced by reciprocally crossing albino animal to heterozygous animal for the albino locus. A total of 32 albino and 59 heterozygous animals were sequenced, detected for genetic variants, and genotyped for polymorphic sites. Manhattan plot showing –log10 P value of χ2 test across the genome. The y axis represents –log_10_ P value and the x axis represents polymorphism coordinates on each chromosome, which is labeled as red (odd number chromosomes) or blue (even number chromosomes). χ2 test P values were corrected using FDR method across the genome-wide data. The black dashed line represents adjusted P value of 0.05.

Hybrids between albino animal to wild type animal exhibit normal pigmentation pattern, suggesting the genetics variant underlying the albinism is recessive. In order to identify the albino-associated genetic variants, we created a hybrid population by first crossing albino *X. hellerii* with a *xmrk*- introgressed *X. hellerii*, which was established by repetitively backcrossing *xmrk*-bearing interspecies hybrids between *X. maculatus* and *X. hellerii* to *X. hellerii*. The rationale of using *xmrk* introgressed line as parental is to rely on *xmrk* mapping as positive control for genetic mapping. In successive breeding pair setup, we used individuals showing normal body and ocular coloration (albino locus heterozygotes) with *xmrk*-associated melanophore and erythrophore pigmentation pattern enhancement, and homozygous albino *xmrk*-negative individuals as parentals. We generated a total of 91 hybrids, including 32 albino (35%) and 59 wild type (65%; Table 1).

**Table 1.** Genotype and phenotype distribution.

| Genetic Type | Color | Total |
| --- | --- | --- |
| a/+ xmrk + | Red | 39 |
| a/+ xmrk - | Green | 20 |
| a/a xmrk + | Red albino | 20 |
| a/a xmrk - | Albino | 12 |
| **Total Fish** |  | **91** |

### Genetics Mapping of Albinism

Inheritance of the *xmrk* gene leads to expansion of erythrophore and melanophore pigmentation pattern. Therefore, the hybrids inherited the *xmrk* exhibit red coloration on the body side and expanded dorsal fin melanophore pigmentation pattern. Since the *xmrk* introgression is known to be on Chr21, it allows us to test the accuracy of genetics mapping. Genetics mapping of genetic locus underlying the enhanced red pigmentation pattern resulted peaks at the region encoding *xmrk* (Fig. S1). There is a single locus identified that is associated to the albinism (multiple test corrected *p*-value < 0.05; Chr 9: 24,167,249 bp – 24,380,394 bp). Allelic inheritance analyses of the hybrid population within this candidate region showed that all albino individuals are homozygous for variant allele, and all wild type individuals are heterozygous (Table S1).

This candidate region from the genetics mapping include 6 genes: *herc2, oca2*, LOC116725874, *gabrg3, gabra5*, and *gabrb3*. Manual curation of genetics variant analyses showed all albino-associated single nucleotide variants (SNVs) within these genes are intronic (25 of 28), intergenic (2 of 28), or synonymous (1 of 28; Table S1).

### Mutations in *oca2* underlie *Xiphophorus* albinism

RNA-sequencing of wild type and albino individuals showed all genes within the peak region are expressed between albino and wild type individuals. However, we found a large region on exon 16 of *oca2* showing absence of sequencing read mapping for albino individuals (Fig. 3A). We hypothesize that *oca2* has a large structural change, e.g., insertion or deletion.

**Figure 3.**
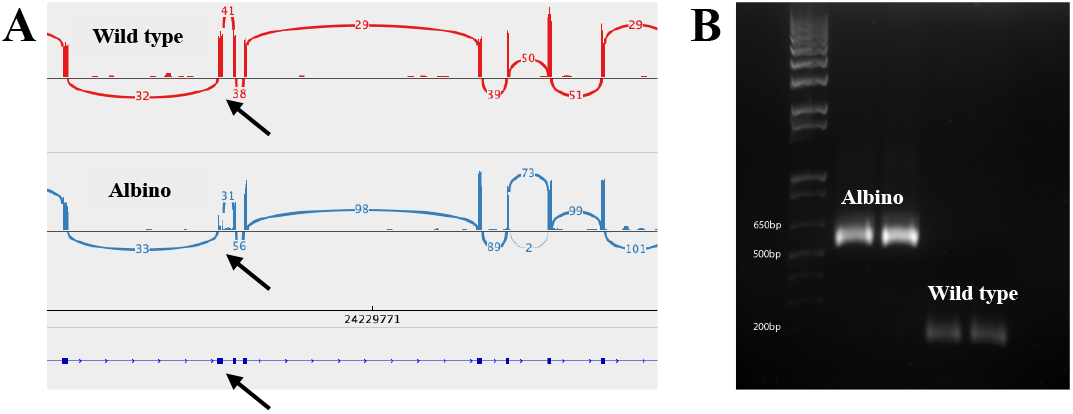
An insertion in *oca2* exon 16 is associated to albinism in *Xiphophorus*. (A) Sashimi plot showing quantification of junction reads connecting exons. The top and middle panel respectively shows quantities of RNA sequencing reads spanning the exon-exon junction (numbers) and mapping locations (density bar graphs) of wild type and albino fish. Bottom panel shows partial *oca2* gene model, with thick lines represented exons. Arrows point to exon 16. (B) Electrophoresis gel images of RT-PCR amplicons representing exon 16 of two albino and two wild type fish.

To test this hypothesis, we sequenced *oca2* exon 16 of both wild type and albino individuals. The results revealed that the albinism-associated oca2 allele contains an approximate 480 bp insertion compared to the wild type allele (Fig. 3B).

### Structural analyses of *oca2* mutation

This insertion introduces a stop codon (Fig. 4A). Structural alignment of the wild type and albino OCA2 proteins (Fig. 4) shows that the mutant allele has a truncation that extend beyond the cytoplasmic region, resulting in the loss of the OCA2 function (Fig. 4C).

**Figure 4.**
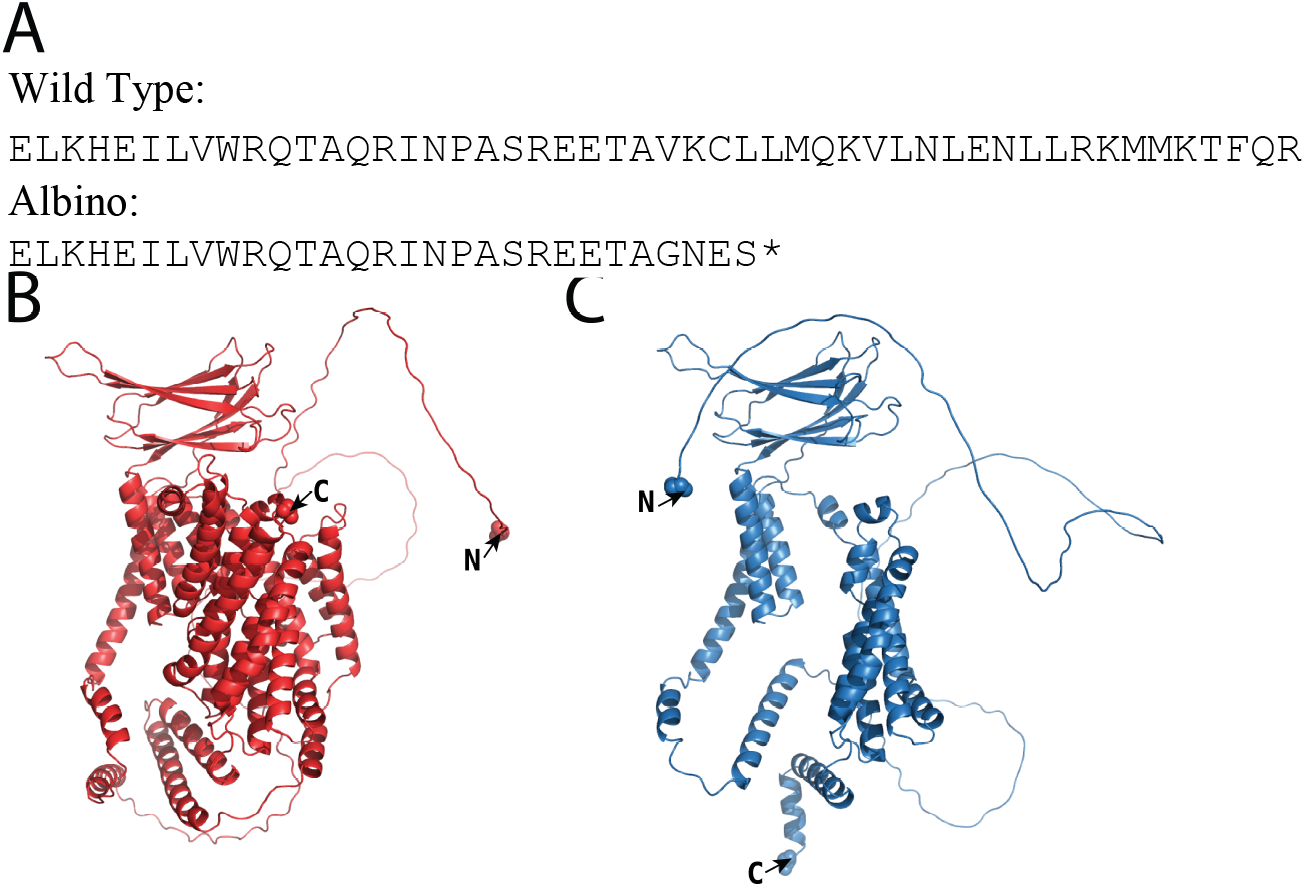
Protein structure analyses of albino allele of *Xiphophorus* OCA2. (A) Peptide sequences wild type and albino OCA2 translated from exon 16 sequences. The mutation in the albino allele introduces an early stop codon, and truncated OCA2 protein. (B) Wild type and (C) mutant protein structures were generated from the Alphafold and the pTM scores are 0.67 and 0.58 respectively.

## Discussion

The gene underlying the OCA2 in human is known as *OCA2* (also known as pink-eyed dilution locus in mice) ^9,32^, which encodes an integral melanosomal protein with 12 predicted transmembrane domains^33,34^. Although the precise function of the OCA2 is not fully understood, it is known to play two role: 1. Melanin biosynthesis: OCA2 is involved in the transport of melanosomal proteins and pH regulation within the melanosome^35^. The melanogenesis pathway relies on TYR activity to initiate melanin synthesis by converting tyrosine to dihydroxyphenylalanine (DOPA) and then to dopaquinone. These reactions occur in an acidic environment, while the following steps of melanogenesis require a neutral pH for TYR to catalyze intermediates that lead to pigment production^36^. OCA2 contributes to a melanosome-specific anion current that modulates melanosomal pH for optimal TYR activity required for melanogenesis^37^. Hence, inhibition of OCA2 functioning is expected to cause changes in melanin synthesis and impact melanosome morphology and quantity^34^. 2. Pigment cell lineage differentiation: The number of pigmented melanosomes decreased in the *Oca2* mutant porcine compared to the wild type pigs^8^. The *Oca2* has been reported to affect proliferation of mouse melanocytes through regulating the expression and activity of melanosomal proteins^38^. The *oca2* knockdown in the wild type pigmented embryos of the surface conspecifics of Mexican cavefish delays the development of pigmented melanophores^6^. The *oca2* mutant in zebrafish shows a reduction in the number of differentiated melanophores and significantly higher numbers of differentiated iridophores than wild type siblings^39^.

In this study, we identified a spontaneous large truncation in the OCA2 protein is associated with albinism in *X. hellerii*. The truncation is expected to affect the protein function significantly by eliminating key structural regions and altering its overall conformation, and may play a role in pigment biosynthesis and/or melanophore differentiation.

The *Xiphophorus* fishes are best recognized for their variations in pigmentation patterns, and elucidating negative genetic interactions underlying spontaneous melanoma development in interspecies hybrids.^21^ In our future studies, we will leverage on the newly found natural *oca2* mutation and investigate *oca2* function in chromophore lineages differentiation and development. In addition, we will cross the albino fish to *Xiphophorus* fish with melanoma driving gene to study how *oca2* affect genetic pathways promoting the melanoma development.

## Acknowledgements

This work was supported by the National Institutes of Health, National Cancer Institute, R15 CA -223964 to Y. Lu, Office of Director R24 OD-031467 to Y. Lu and M. Schartl, R2R1 accelerator award from Texas State University to Y. Lu and M. Schartl.

## Supplemental Figure

**Figure S1. Genetic mapping of *xmrk***

A total of 59 animals exhibiting red coloration enhancement and 32 animals exhibit wild type erythrophore pigmentation pattern were sequenced, detected for genetic variants, and genotyped for polymorphic sites. Manhattan plot showing –log10 P value of χ2 test across the genome. The y axis represents –log_10_ P value and the x axis represents polymorphism coordinates on each chromosome, which is labeled as red (odd number chromosomes) or blue (even number chromosomes).

## Supplemental Table

Table S1. Genetic Variants in Albino vs. Wild type

