## Supplemental figure and table for "A Recessive *oca2* Mutation Underlies Albinism in *Xiphophorus* fish"

| loci | Albino het | Albino homo | WT het | WT homo | p.value | Gene | Exon |
|---|---|---|---|---|---|---|---|
| 24167249 | 0 | 29 | 52 | 0 | 5.30E-23 | herc2 | intron |
| 24181019 | 0 | 30 | 53 | 0 | 1.21E-23 | herc2 | intron |
| 24181791 | 0 | 29 | 56 | 0 | 7.69E-24 | herc2 | intron |
| 24200393 | 0 | 27 | 53 | 0 | 2.15E-22 | herc2 | intron |
| 24209457 | 0 | 28 | 53 | 0 | 8.79E-23 | oca2 | intron |
| 24218279 | 0 | 28 | 53 | 0 | 8.79E-23 | oca2 | intron |
| 24219340 | 0 | 27 | 52 | 0 | 3.63E-22 | oca2 | intron |
| 24223691 | 0 | 26 | 56 | 0 | 8.15E-23 | oca2 | intron |
| 24227175 | 0 | 29 | 55 | 0 | 1.28E-23 | oca2 | intron |
| 24229081 | 0 | 27 | 50 | 0 | 9.80E-22 | oca2 | intron |
| 24236420 | 0 | 32 | 52 | 0 | 1.86E-24 | oca2 | intron |
| 24238679 | 0 | 29 | 58 | 0 | 2.63E-24 | XR_004340503.1 (oca2 intron) | exon1 (synonymous) |
| 24246268 | 0 | 31 | 51 | 0 | 8.68E-24 | oca2 | intron |
| 24254931 | 0 | 30 | 53 | 0 | 1.21E-23 | intergenic | |
| 24280454 | 0 | 28 | 53 | 0 | 8.79E-23 | gabrg3 | intron |
| 24284424 | 0 | 30 | 53 | 0 | 1.21E-23 | gabrg3 | intron |
| 24290340 | 0 | 28 | 53 | 0 | 8.79E-23 | gabrg3 | intron |
| 24302907 | 0 | 28 | 50 | 0 | 3.60E-22 | gabrg3 | intron |
| 24310161 | 0 | 29 | 53 | 0 | 3.37E-23 | gabrg3 | intron |
| 24313969 | 0 | 31 | 51 | 0 | 8.68E-24 | intergenic | |
| 24318741 | 0 | 30 | 50 | 0 | 4.02E-23 | gabra5 | intron |
| 24318752 | 0 | 29 | 48 | 0 | 2.67E-22 | gabra5 | intron |
| 24335105 | 0 | 27 | 52 | 0 | 3.63E-22 | gabra5 | intron |
| 24366765 | 0 | 31 | 49 | 0 | 1.70E-23 | gabrb3 | intron |
| 24367799 | 0 | 27 | 55 | 0 | 7.11E-23 | gabrb3 | intron |
| 24368808 | 0 | 28 | 53 | 0 | 8.79E-23 | gabrb3 | intron |
| 24379271 | 0 | 27 | 55 | 0 | 7.11E-23 | gabrb3 | intron |
| 24380394 | 0 | 31 | 50 | 0 | 1.23E-23 | gabrb3 | intron |

**Table S1**

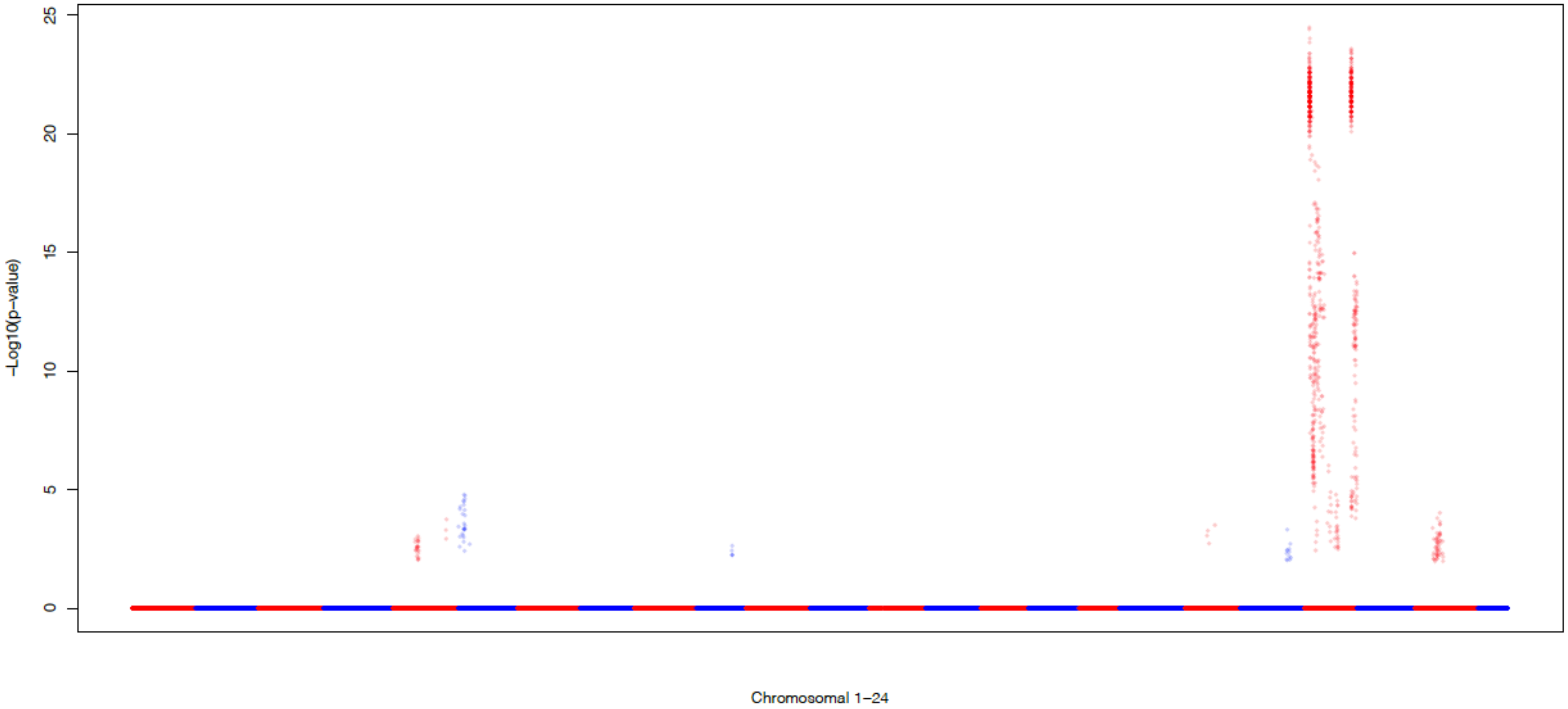


**Figure S1**
