## Supplementary material for "A Recessive *oca2* Mutation Underlies Albinism in *Xiphophorus* fish": Supplental data

>XM_032572273.1 PREDICTED: Xiphophorus hellerii OCA2 melanosomal transmembrane protein (oca2), mRNA
TCTTTCGCTGGGGAGAGCGCAGGAGAGAAGAAACTCCTCACCTTCTCTTGTTTTCAACCACGCCTCTGTG
CTCCGACGTTACTGCAGCCTCCTCATCCCACCCCCAGCAAAAAAGAAGAAGAAAAAAAAAAGTATTAATCA
AACAAAAAGTTGAAAATAATTTGGATTACTGAAAAGTAAAGTTTTGCAGCAGATTTTTTTTTTTAATTCA
GATCCAACACATCGTTCCTACCTGCGCCTTCAACATCATCCAGAGCGACTTTTTTTTTTTGTGCATCTCT
CAAGGCCTCAGAATGTATCTGGAGAACAAAAGCAACCTGGAGATTGAAATGTCTCAGACTTCGGCCGCGG
GCCATCAGGGCAGGAGCGGCCTCAGGTCGCCGGAGGCCGTGCCGGGGAACTTTGGGGAGCTGCGTCTGCT
TCAGGGGATCTCAGAGAGAGGAGACACCCAGAAGCTGCAGCCTGCAGGACAGTGCGTCATCCATGCTGAG
GACCTGGTTCCGGTTCGGCAAGAGAACGGTTTCAGGAACAACTTCAACGCTGTCAACTTTCTCAGAGGAA
AAGCCAATTCAGCCAACCTTACAGAACGGTCGCCTCTGCTGCGATTCTCCCAAGATGACAGTATTACGTA
CATGACAGTACATGAGCCCAGCTTCGTTGGAGTAGAGGAGCCATGGGACAACTGCTGCATGGACACAGAG
AGACGTTACCGCCTGGGCAGCGAGGTGGCCGGCCTGTCGCGCTCCAACTCCTCAGAAAAGTACGAGACAC
TGGAAAAACCTGAACCGCGGCTTCAGACTCACCAGCAGTATAAAATGCTGCCTCAGAGTGTCCAAGGTTGT
GGCTATGTTTGCGATTGTTGTTCTCTCCAGTCTGTTCTTCAGTATGTACCCGGATCGGGACAACCCCTGG
AGGATGCTGGCCGTCTCTCCCACAGACAGCTTCTCCATGAACGTGACAGACTTTCGGGATAACGCCTTGC
TGAAGTTACAAGTGGCGGGGCCTTTTGCAGGAAGGGCTGTGGACTCCGCCTCCCAGGAGTTCATCCTGAT
CCAGGTGGAGCAGGCCGAGGAGACGGGGCAGCGGAGGAGGAGGACTCAGCAGGTGACTCATAACTGGACC
ATTCCCCTCCATCCTGAACGGAGTGACCAGAAGGTGTCAACAAGAACCTTTGAGATGCTCAGCAGTGACC
CCATCGTCATCACGATTCAGGCCTCCCTGCAGGATAACGACGTGGTGCCTCTGTCCATGACCCACCAATC
GCTCTACGTTACGAAGGAGACGCAGGTGGCTATAGCGGGAATCATCCTGGCCGGGGTCTATGTTCTCATC
ATCTTTGAGATTGTGCATCGCACTCTGGCTGCCATGTTGGGATCCTTAGCAGCTTTAGCAGCCCTGGCTA
TCATTGGTGACAGACCCAGCCTGGTCACTGTGGTGGAGTGGATCGACTACGAGACGCTGGCTCTGCTGTT
TGGCATGATGGTGCTGGTGGCCATTTTTTCAGAGACAGGCTTCTTTGATTATTGTGCGGTGAAGGCCTAC
CAGTTATCCAGAGGGAGGGTGTGGCCTATGATCATCATCCTGTGTCTGATTGCTGCCATTCTGTCTGCCT
TCCTGGATAATGTGACCACCATGATGCTTTTCACTCCTGTCACCATCAGGTTGTGTGAGGTGCTTAATCT
AGACCCCAGACATGTCCTGATTGCAGAAGTAATCTTCACCAACATCGGAGGGGCCGCCACAGCTGTCGGA
GATCCACCCAACGTGATTATCGTGTCCAACCAGGGCCTGCGCAGAGAAGGAATAGACTTTGCCAGCTTCA
CAGGCTACATGTTTCTTGGAATATGCCTCGTCCTCCTCACCTCATTCCCTTTCCTCAGGATGCTCTACTG
GAACAAAAAACTTTACAACAAAGAGTCCATTGAAATAGTTGAACTGAAGCATGAGATCCTGGTCTGGAGG
CAAACGGCTCAGCGCATTAACCCGGCCAGCCGGGAGGAGACAGCTGTGAAATGCCTTCTGATGCAAAAAG
TGCTCAACCTGGAAAATCTGCTCCGCAAGATGATGAAAACCTTCCAGAGACAAATTTCTCAGGAGGATAA
AAACTGGGAACAAAACATTCAGGAGCTGCAGAAGAAGCACAGAATAACAGACACGGTTCTGTTGGTGAAA
TGTGCCTCCGTCCTTGGTGTGGTGATCTTCATGTTCTTTCTCAACTCCTTTGTTCCCAGCATTCATCTGG
ACCTGGGTTGGATCGCCATCCTGGGGGCCCTCTGGCTCCTAGTTCTGGCCGATGTCCAGGACTTTGAGAT
CATCCTGCACCGGGTGGAGTGGGCCACTCTGCTGTTCTTCGCGGGCCTCTTTGTTTTAATGGAGGCTTTA
GCCCAGCTGCAGCTTATCGACTATATTGGAGAGCAGACGGCGCTGCTGATAAAAGCGGTGCCAGAGGACC
AGCGCCTGGCTGTGGCCATTATCTTGGTGTTGTGGGTCTCCGCTCTGGCTTCTTCTCTCATCGACAACAT
TCCCTTCACTGCCACCATGATCCCAGTTCTCATCAACCTGAGTCAGGATGTGGACGTCAACCTCCCTGTG
AAGCCTCTCATCTTTGCCTTGGCTATGGGGGCCTGCCTCGGTGGAAACGGCACGTTGATCGGGGCGTCAG
CCAACGTCGTGTGTGCAGGCATCGCCGAGCAACATGGATACGGTTTTTCCTTCGTGGAGTTCTTCAAGCT
GGGTTTCCCCATGATGGTGATGACGTGCCTGGTCGCCATGTGCTACCTGCTGGCCACACACATCGGCCTG
GGATGGAACATGTGAACCGCGCCGGCCAAAGAGACGGAGTGCGTTGGCACACGTGCGTGTACGTTTACGT
ACGGGCGTGTATGCGTGTGTGTAGCCACTTCTCTAGGAGTTGCAGCTCACTGTAAATGTGATGTGTAAGC
TGACCGACCAATCAGATTACAAGCAGAACATGAAGCACTGCAGCCATGTTTTTTTTGTTTGTTTGTTTGT
TTGTTTTTTTTTAAGCGTGGGGGATGTGTGGACCATCAGATTTACAAGAGTTTTGAACAAAAGCCGACTC
TGAGAGAAGCGTTTCTTGGCTGCACTTCATATCTGTGTTCGTGGTGCTACGGCGTCAGAGGGAACAGCAG
CTGCCGTCTGCCGTCTCTCCTTTGGAAATCCGTTTACAAAAGACGTCAAATGATTTCTTCAATCACTTAT
TGTAACCCTTTAATGAAATATACAACTAGAGACTGCTGATTGATTGTTTTGTACCAATTGAAACATCTGA
AGAAAGGCCCAGTGAAAGGGATTTTATACCAGTTTATGTCTGTATTTCACTGTTACGATATACGAAATTT
TCAAAGCAATAATTTGTTTTTCAGATTTCTCTGATTGTGTCCGGTCATTTTAATTGTACTGATTGGTCCC
AGACTGGGTCTGTCACTCTGTCAAGCTCTGGTTGTTTTCTGTATTTCTTGTACGGCTCAACAACACGGAA
CGTCTTCTGTGTTCTCCGAATAAACGTTTCAGAAGACGAATAATAAATGGTGTCATATCTCTGGA

>Albino oca2, mRNA
TCTTTCGCTGGGGAGAGCGCAGGAGAGAAGAAACTCCTCACCTTCTCTTGTTTTCAACCACGCCTCTGTG
CTCCGACGTTACTGCAGCCTCCTCATCCCACCCCCAGCAAAAAAGAAGAAGAAAAAAAAAGTATTAATCA
AACAAAAAGTTGAAAATAATTTGGATTACTGAAAAGTAAAGTTTTGCAGCAGATTTTTTTTTTTAATTCA
GATCCAACACATCGTTCCTACCTGCGCCTTCAACATCATCCAGAGCGACTTTTTTTTTTTGTGCATCTCT
CAAGGCCTCAGAATGTATCTGGAGAACAAAAGCAACCTGGAGATTGAAATGTCTCAGACTTCGGCCGCGG
GCCATCAGGGCAGGAGCGGCCTCAGGTCGCCGGAGGCCGTGCCGGGGAACTTTGGGGAGCTGCGTCTGCT
TCAGGGGATCTCAGAGAGAGGAGACACCCAGAAGCTGCAGCCTGCAGGACAGTGCGTCATCCATGCTGAG
GACCTGGTTCCGGTTCGGCAAGAGAACGGTTTCAGGAACAACTTCAACGCTGTCAACTTTCTCAGAGGAA
AAGCCAATTCAGCCAACCTTACAGAACGGTCGCCTCTGCTGCGATTCTCCCAAGATGACAGTATTACGTA
CATGACAGTACATGAGCCCAGCTTCGTTGGAGTAGAGGAGCCATGGGACAACTGCTGCATGGACACAGAG
AGACGTTACCGCCTGGGCAGCGAGGTGGCCGGCCTGTCGCGCTCCAACTCCTCAGAAAAGTACGAGACAC
TGGAAAACCTGAACCGCGGCTTCAGACTCACCAGCAGTATAAAATGCTGCCTCAGAGTGTCCAAGGTTGT
GGCTATGTTTGCGATTGTTGTTCTCTCCAGTCTGTTCTTCAGTATGTACCCGGATCGGGACAACCCCTGG
AGGATGCTGGCCGTCTCTCCCACAGACAGCTTCTCCATGAACGTGACAGACTTTCGGGATAACGCCTTGC
TGAAGTTACAAGTGGCGGGGCCTTTTGCAGGAAGGGCTGTGGACTCCGCCTCCCAGGAGTTCATCCTGAT
CCAGGTGGAGCAGGCCGAGGAGACGGGGCAGCGGAGGAGGAGGACTCAGCAGGTGACTCATAACTGGACC
ATTCCCCTCCATCCTGAACGGAGTGACCAGAAGGTGTCAACAAGAACCTTTGAGATGCTCAGCAGTGACC
CCATCGTCATCACGATTCAGGCCTCCCTGCAGGATAACGACGTGGTGCCTCTGTCCATGACCCACCAATC
GCTCTACGTTACGAAGGAGACGCAGGTGGCTATAGCGGGAATCATCCTGGCCGGGGTCTATGTTCTCATC
ATCTTTGAGATTGTGCATCGCACTCTGGCTGCCATGTTGGGATCCTTAGCAGCTTTAGCAGCCCTGGCTA
TCATTGGTGACAGACCCAGCCTGGTCACTGTGGTGGAGTGGATCGACTACGAGACGCTGGCTCTGCTGTT
TGGCATGATGGTGCTGGTGGCCATTTTTTCAGAGACAGGCTTCTTTGATTATTGTGCGGTGAAGGCCTAC
CAGTTATCCAGAGGGAGGGTGTGGCCTATGATCATCATCCTGTGTCTGATTGCTGCCATTCTGTCTGCCT
TCCTGGATAATGTGACCACCATGATGCTTTTCACTCCTGTCACCATCAGGTTGTGTGAGGTGCTTAATCT
AGACCCCAGACATGTCCTGATTGCAGAAGTAATCTTCACCAACATCGGAGGGGCCGCCACAGCTGTCGGA
GATCCACCCAACGTGATTATCGTGTCCAACCAGGGCCTGCGCAGAGAAGGAATAGACTTTGCCAGCTTCA
CAGGCTACATGTTTCTTGGAATATGCCTCGTCCTCCTCACCTCATTCCCTTTCCTCAGGATGCTCTACTG
GAACAAAAAACTTTACAACAAAGAGTCCATTGAAATAGTTGAACTGAAGCATGAGATCCTGGTCTGGAGG
CAAACGGCTCAGCGCATTAACCCGGCCAGCCGGGAGGAGACAGCTGGAAATGAATCC**TAA**GCCCAAATTC
GAACTGGAAGACTCCTTAGCCCCACCTGAATCATCCAATCGGCGCGAGACGCTCATCACCACCGACCTAT
CAACGCTGGACCAAGCCACGTCACGTCCAATCACCGACCAAGGCCCAGTTGATAAGGACCTCCATCCATG
GCATCTTCCGCTTTCCAGCTGCGCTCAACCCTCCTCCACCCCTTCTCCACCTCCACTCTTCAGGCTCCCA
TCCTTCACCGACCTCCACCACCAGTGTCGGGTCCCGGGGGATGACATTCCTAACCTACATCGGGAATGCT
CATCCGAGCTGATCGCAATTCACAGCTCAATGTCCTCAGCCGGGGCCCGAGCGCCAAAGAACTTTAAATT
CATGCATGTCAATAAACCTTTTTAATCGCTGTTTTTCGTCTGAGTCTCTCCAGATTGAAATGCCTTCTGA
TGCAAAAAGTGCTCAACCTGGAAAATCTGCTCCGCAAGATGATGAAAACCTTCCAGAGACAAATTTCTCA
GGAGGATAAAAACTGGGAACAAAACATTCAGGAGCTGCAGAAGAAGCACAGAATAACAGACACGGTTCTG
TTGGTGAAATGTGCCTCCGTCCTTGGTGTGGTGATCTTCATGTTCTTTCTCAACTCCTTTGTTCCCAGCA
TTCATCTGGACCTGGGTTGGATCGCCATCCTGGGGGCCCTCTGGCTCCTAGTTCTGGCCGATGTCCAGGA
CTTTGAGATCATCCTGCACCGGGTGGAGTGGGCCACTCTGCTGTTCTTCGCGGGCCTCTTTGTTTTAATG
GAGGCTTTAGCCCAGCTGCAGCTTATCGACTATATTGGAGAGCAGACGGCGCTGCTGATAAAAGCGGTGC
CAGAGGACCAGCGCCTGGCTGTGGCCATTATCTTGGTGTTGTGGGTCTCCGCTCTGGCTTCTTCTCTCAT
CGACAACATTCCCTTCACTGCCACCATGATCCCAGTTCTCATCAACCTGAGTCAGGATGTGGACGTCAAC
CTCCCTGTGAAGCCTCTCATCTTTGCCTTGGCTATGGGGGCCTGCCTCGGTGGAAACGGCACGTTGATCG
GGGCGTCAGCCAACGTCGTGTGTGCAGGCATCGCCGAGCAACATGGATACGGTTTTTCCTTCGTGGAGTT
CTTCAAGCTGGGTTTCCCCATGATGGTGATGACGTGCCTGGTCGCCATGTGCTACCTGCTGGCCACACAC
ATCGGCCTGGGATGGAACATGTGAACCGCGCCGGCCAAAGAGACGGAGTGCGTTGGCACACGTGCGTGTA
CGTTTACGTACGGGCGTGTATGCGTGTGTGTAGCCACTTCTCTAGGAGTTGCAGCTCACTGTAAATGTGA
TGTGTAAGCTGACCGACCAATCAGATTACAAGCAGAACATGAAGCACTGCAGCCATGTTTTTTTTGTTTG
TTTGTTTGTTTGTTTTTTTTTAAGCGTGGGGGATGTGTGGACCATCAGATTTACAAGAGTTTTGAACAAA
AGCCGACTCTGAGAGAAGCGTTTCTTGGCTGCACTTCATATCTGTGTTCGTGGTGCTACGGCGTCAGAGG
GAACAGCAGCTGCCGTCTGCCGTCTCTCCTTTGGAAATCCGTTTACAAAAGACGTCAAATGATTTCTTCA
ATCACTTATTGTAACCCTTTAATGAAATATACAACTAGAGACTGCTGATTGATTGTTTTGTACCAATTGA
AACATCTGAAGAAAGGCCCAGTGAAAGGGATTTTATACCAGTTTATGTCTGTATTTCACTGTTACGATAT
ACGAAATTTTCAAAGCAATAATTTGTTTTTCAGATTTCTCTGATTGTGTCCGGTCATTTTAATTGTACTG
ATTGGTCCCAGACTGGGTCTGTCACTCTGTCAAGCTCTGGTTGTTTTCTGTATTTCTTGTACGGCTCAAC

AACACGGAACGTCTTCTGTGTTCTCCGAATAAACGTTTCAGAAGACGAATAATAAATGGTGTCATATCTC
TGGA

Stop Codon: Bold & Underlined
Insertion: Underlined

>XP_032428164.1 OCA2 [Xiphophorus hellerii]
MYLENKSNLEIEMSQTSAAGHQGRSGLRSPEAVPGNFGELRLLQGISERGDTQKLQPAGQCVIHAEDLVP
VRQENGFRNNFNAVNFLRGKANSANLTERSPLLRFSQDDSITYMTVHEPSFVGVEEPWDNCCMDTERRYR
LGSEVAGLSRSNSSEKYETLENLNRGFRLTSSIKCCLRVSKVVAMFAIVVLSSLFFSMYPDRDNPWRMLA
VSPTDSFSMNVTDFRDNALLKLQVAGPFAGRAVDSASQEFILIQVEQAEETGQRRRRTQQVTHNWTIPLH
PERSDQKVSTRTFEMLSSDPIVITIQASLQDNDVVPLSMTHQSLYVTKETQVAIAGIILAGVYVLIIFEI
VHRTLAAMLGSLAALAALAIIGDRPSLVTVVEWIDYETLALLFGMMVLVAIFSETGFFDYCAVKAYQLSR
GRVWPMIIILCLIAAILSAFLDNVTTMMLFTPVTIRLCEVLNLDPRHVLIAEVIFTNIGGAATAVGDPPN
VIIVSNQGLRREGIDFASFTGYMFLGICLVLLTSFPFLRMLYWNKKLYNKESIEIVELKHEILVWRQTAQ
RINPASREETAVKCLLMQKVLNLENLLRKMMKTFQRQISQEDKNWEQNIQELQKKHRITDTVLLVKCASV
LGVVIFMFFLNSFVPSIHLDLGWIAILGALWLLVLADVQDFEIILHRVEWATLLFFAGLFVLMEALAQLQ
LIDYIGEQTALLIKAVPEDQRLAVAIILVLWVSALASSLIDNIPFTATMIPVLINLSQDVDVNLPVKPLI
FALAMGACLGGNGTLIGASANVVCAGIAEQHGYGFSFVEFFKLGFPMMVMTCLVAMCYLLATHIGLGWNM

>Albino OCA2
MYLENKSNLEIEMSQTSAAGHQGRSGLRSPEAVPGNFGELRLLQGISERGDTQKLQPAGQCVIHAEDLVP
VRQENGFRNNFNAVNFLRGKANSANLTERSPLLRFSQDDSITYMTVHEPSFVGVEEPWDNCCMDTERRYR
LGSEVAGLSRSNSSEKYETLENLNRGFRLTSSIKCCLRVSKVVAMFAIVVLSSLFFSMYPDRDNPWRMLA
VSPTDSFSMNVTDFRDNALLKLQVAGPFAGRAVDSASQEFILIQVEQAEETGQRRRRTQQVTHNWTIPLH
PERSDQKVSTRTFEMLSSDPIVITIQASLQDNDVVPLSMTHQSLYVTKETQVAIAGIILAGVYVLIIFEI
VHRTLAAMLGSLAALAALAIIGDRPSLVTVVEWIDYETLALLFGMMVLVAIFSETGFFDYCAVKAYQLSR
GRVWPMIIILCLIAAILSAFLDNVTTMMLFTPVTIRLCEVLNLDPRHVLIAEVIFTNIGGAATAVGDPPN
VIIVSNQGLRREGIDFASFTGYMFLGICLVLLTSFPFLRMLYWNKKLYNKESIEIVELKHEILVWRQTAQ
RINPASREETA<u>GNES</u>

Mutation: Underlined
